# Generation and analysis of context-specific genome-scale metabolic models derived from single-cell RNA-Seq data

**DOI:** 10.1101/2022.04.25.489379

**Authors:** Johan Gustafsson, Jonathan L. Robinson, Fariba Roshanzamir, Rebecka Jörnsten, Eduard J Kerkhoven, Jens Nielsen

**Affiliations:** Department of Biology and Biological Engineering, Chalmers University of Technology, Gothenburg, Sweden; Wallenberg Center for Protein Research, Chalmers University of Technology, Gothenburg, Sweden; BioInnovation Institute, Copenhagen, Denmark; Mathematical Sciences, University of Gothenburg and Chalmers University of Technology, Gothenburg, Sweden

## Abstract

Single-cell RNA sequencing has the potential to unravel the differences in metabolism across cell types and cell states in both the healthy and diseased human body. The use of existing knowledge in the form of genome-scale metabolic models (GEMs) holds promise to strengthen such analyses, but the combined use of these two methods requires new computational methods. Here, we present a method for generating cell-type-specific genome-scale models from clusters of single-cell RNA-Seq profiles. Specifically, we developed a method to estimate the number of cells required to pool to obtain stable models, a bootstrapping strategy for estimating statistical inference, and a faster version of the tINIT algorithm for generating context-specific GEMs. In addition, we evaluated the effect of different RNA-Seq normalization methods on model topology and differences in models generated from single-cell and bulk RNA-Seq data. We applied our methods on data from mouse cortex neurons and cells from the tumor microenvironment of lung cancer and in both cases found that almost every cell subtype had a unique metabolic profile, emphasizing the need to study them separately rather than to build models from bulk RNA-Seq data. In addition, our approach was able to detect cancer-associated metabolic differences between cancer cells and healthy cells, showcasing its utility. With the ever-increasing availability of single-cell RNA-Seq datasets and continuously improved GEMs, their combination holds promise to become an important approach in the study of human metabolism.

## Introduction

Genome-scale metabolic models (GEMs) have been extensively used to further our understanding of metabolism in both unicellular organisms such as yeast and bacteria (1,2), and multicellular species such as humans (3–5). For multicellular species, the existence of many different cell types and tissues poses a challenge for metabolic modeling since the full reaction network encoded by the genome is typically not present in such tissues or cell types. To remedy this, several methods have been developed that utilize RNA sequencing data or proteomics to detect the active subnetwork in a sample (6–8), such as the task-driven integrative network inference for tissues (tINIT) algorithm. Such methods start with a full model and generate context-specific models, containing only the active portion of the network within a given tissue or cell type.

Each tissue in the human body contains many cell types and cell subtypes, where each of these often have several transcriptional states. Bulk RNA-Seq measurements are useful for generating context-specific models that describe the collective metabolism of the cell types in a tissue and if used with fluorescence-activated cell sorting (FACS), for example, can be used to target individual cell types. However, the technique is limited to cell types and states that can be separated by cell surface markers, which must be decided beforehand. Availability of single-cell RNA-Seq (scRNA-Seq) presents a new opportunity to generate context-specific models at the level of individual cell types and cell states.

Obtaining a representative gene expression profile for a cell type can be challenging due to technical variation in the data (9,10), where data sparsity in particular is a substantial challenge when generating context-specific models from single-cell RNA-Seq data. The variation in the data, particularly in single-cell data from droplet-based methods, is dominated by sampling effects, often requiring averaging (pooling) the individual profiles of thousands of cells to obtain the same expected variation as observed between bulk RNA-Seq samples (11). Previously reported methods for generating context-specific models for single cells either focus on small simplified models targeting highly expressed enzymes(12) or use different strategies to integrate data from neighboring cells(13), also focusing on highly expressed pathways. While these methods are useful for finding differences in metabolism, they do not focus on capturing the entire metabolic network of a cell type, with the purpose of using these networks for further simulation. Others have generated context-specific models from pooled single-cell RNA-Seq data (14,15), but do not fully address the statistical uncertainties introduced by the data sparsity.

In this work, we developed methods for generating context-specific GEMs from pools (often clusters) of single-cell RNA-Seq profiles. The methods include an estimation of the required pool size and a bootstrapping strategy to estimate uncertainties in the ensuing reaction subnetwork. Since the bootstrapping strategy requires the generation of many models, we developed a new optimized version of tINIT called ftINIT (fast tINIT), which is substantially faster than the previous versions. We applied our methods on a mouse brain single-cell RNA-Seq dataset, showcasing the ability of the methods to identify differences in metabolic capabilities across neurons. Furthermore, we used our methods to investigate a dataset from the tumor microenvironment of lung cancer and found unique metabolic capabilities of cells known to be associated with cancer.

## Results

### Generation of cell-type-specific models

To investigate the difference in active metabolic network between cell types, we generated context-specific genome-scale models by reducing the generic GEM Human1 (3) based on scRNA-Seq data (Fig. 1A). The process starts with generating clusters of single cells by cell type. To enable comparison across cell types, it is desirable to estimate the uncertainty in modeling results and apply statistical inference. For scRNA-Seq data, we propose to generate GEMs from multiple bootstraps of cells from each cluster to assess the robustness of the modeling results, since the total number of UMIs/reads is usually too small for applying statistics across models generated from separate biological samples (Note S1). Each bootstrap sample is then pooled into an RNA-Seq profile, and context-specific models are generated for each such sample. Further analyses, such as the evaluation of metabolic tasks, is performed for each individual model and statistical methods can be applied across cell types, where each cell type is represented by a group of bootstrap models.

**Fig. 1:**
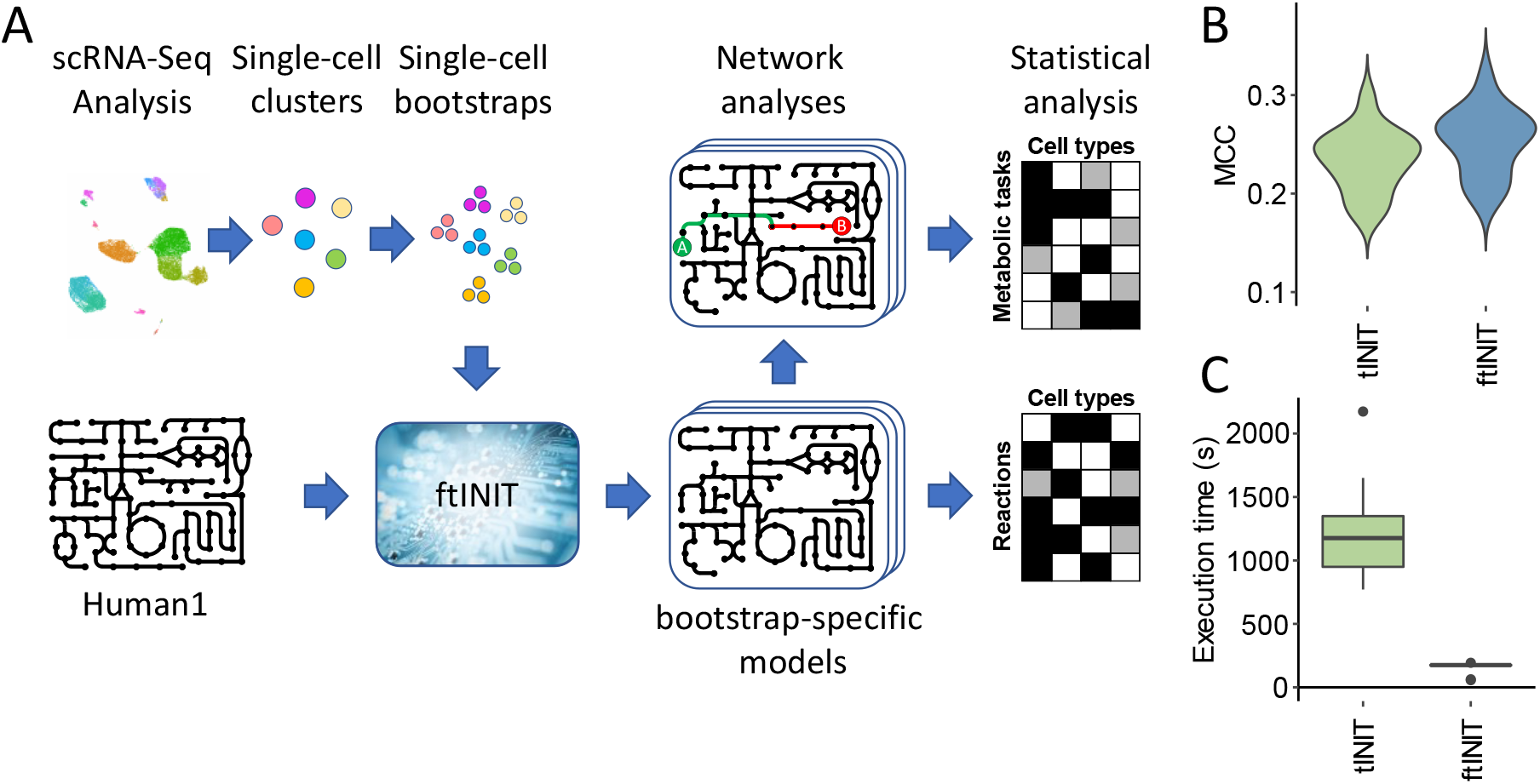
Generation of context-specific models from single-cell RNA-Seq data. A. Overview of model generation and analysis. Cells are first clustered in the single-cell RNA-Seq data. Bootstraps of single cells are then generated from each cluster, followed by pooling of the single cells to form a transcriptomic profile, which together with the template model Human1 is used as input to ftINIT to generate context-specific models for each bootstrap of each cell type. Network analyses in the form of metabolic task analysis is then performed per bootstrap model, and statistical analysis is applied across bootstraps to decide if a reaction or metabolic task is available, unavailable, or uncertain in each cell type. B. Evaluation of the ability to predict essential genes by the models created using tINIT and ftINIT, respectively, for 15 cell lines from DepMap. The performance is measured using the Matthews correlation coefficient (MCC). C. Execution times for tINIT and ftINIT applied on 10 samples from GTEx.

The tINIT (8) algorithm has previously been developed for generating context-specific GEMs based on either transcriptomic or proteomic data. A drawback of the method is the computation time, which for more complex models such as Human1 can range from 15 minutes to 3 hours on a standard laptop computer for a single tissue model. Since the bootstrapping strategy requires generation of large quantities of context-specific GEMs, we sought to optimize the method, resulting in ftINIT (Note S2). The results from ftINIT are different from that of previous versions; ftINIT for example employs a different strategy for reactions lacking gene associations, where many such reactions are included rather than excluded, and the ftINIT optimization is divided into several steps to reduce computation time. We evaluated the performance of the previous and new version of tINIT using gene essentiality analysis on 15 cell lines from DepMap (16,17), which showed a similar ability of the produced models to predict gene essentiality (Fig. 1B). We compared ftINIT with the previous version, and models generated from transcriptomic data from the GTEx project (18) grouped per tissue in a comparable way as the original tINIT method (Fig S1-S2), although with slightly more spread. We likewise evaluated the reduction in execution time, which was substantial (Fig. 1C).

### Technical evaluation of modeling from single-cell RNA-Seq data

To evaluate the technical limitations of single-cell RNA-Seq data, we first investigated the reproducibility of context-specific GEMs generated from such data. Specifically, we compared models generated from non-overlapping randomly selected cell subpopulations from the same cell type cluster (Fig. 2A). Surprisingly, thousands of cells were typically needed for droplet-based single-cell data to generate models with the same variation as between bulk samples, and increasing the pool size beyond 10,000 cells continues to improve the stability of the cell-specific GEMs. Furthermore, the number of cells required for stable model generation varied across datasets, where datasets with more UMIs per cell (“HCA CB T” in Fig. 2) generally required fewer cells, emphasizing the need to evaluate the required pool size per dataset. Direct model comparison, as shown in Fig. 2A, is impractical due to the large computational cost required for such a method. We therefore investigated the use of our previously developed single-cell variation estimation method DSAVE (11) to quantify the variation between pools of cells, which takes less than a minute to run on a standard laptop computer. DSAVE demonstrated reasonable agreement in the estimated required pool size (Fig. 2B). Based on our results, when generating context-specific models we recommend pooling at least the number of cells required to reach the DSAVE total variation score of the bulk reference.

**Fig. 2:**
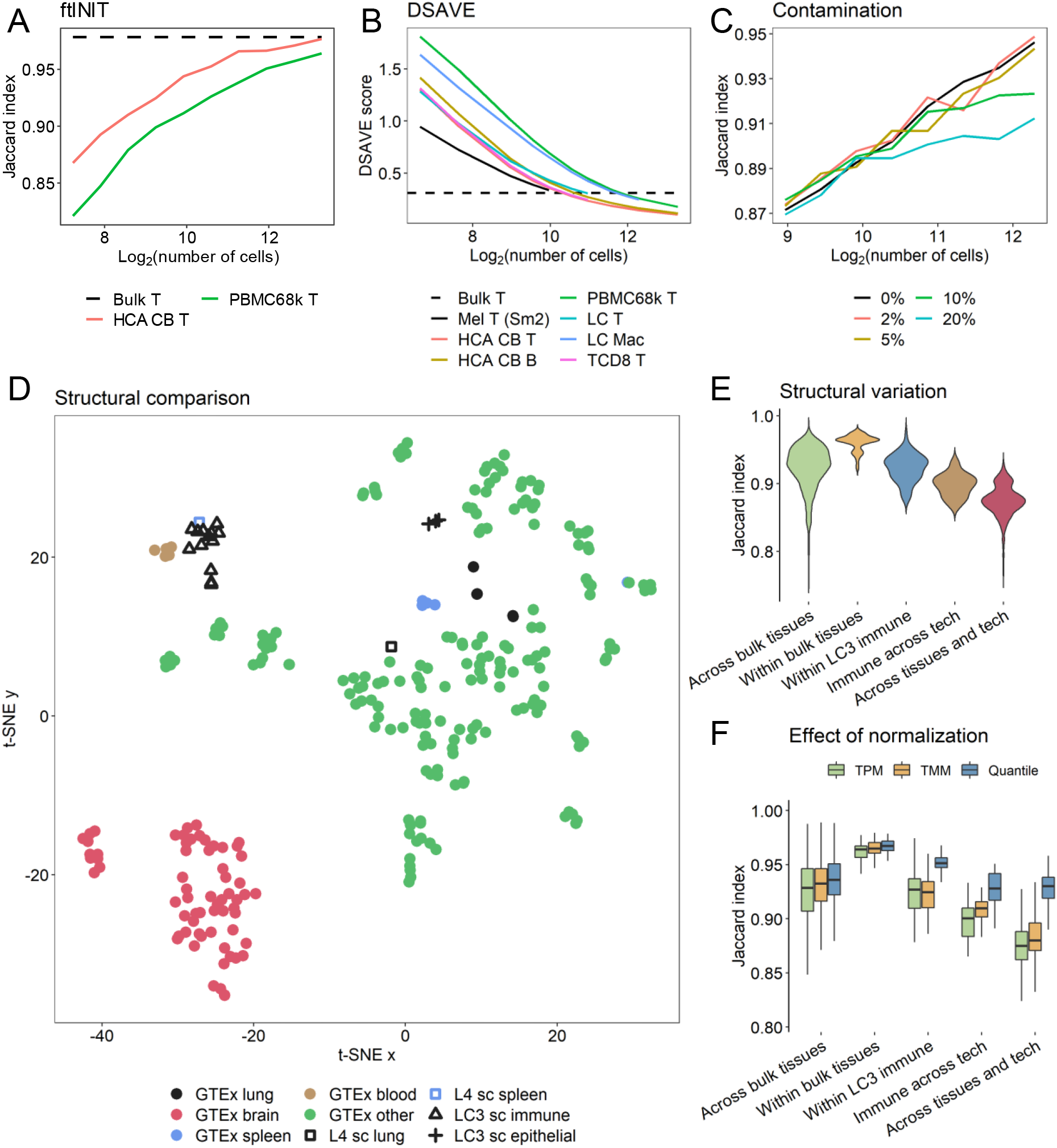
Technical evaluation of generating context-specific models from scRNA-Seq data. A. Reproducibility of context-specific model generation per single-cell pool size, using ftINIT. For each comparison, two non-overlapping sets of single cells were pooled from the same cell type (T cells) and dataset, followed by GEM generation by ftINIT. The two models were compared based on their reaction content. The data shown is an average of 30 such repetitions per pool size. The bulk reference value represents a comparison of bulk T cells (FACS-sorted). B. DSAVE total variation score per pool size. C. The effect of contaminating cell pools with cells of another cell type. T cell populations from the LC dataset were contaminated to a varying degree with tumor cells from the same dataset and patient and compared to the pure T cell population. Reaction scores were calculated per gene (Methods) and converted to on/off with a threshold of 1 CPM, followed by direct comparison without involving ftINIT. D. Structural comparison of models generated by ftINIT from various sources, both GTEx bulk samples from 53 tissue types and different single-cell datasets. Two samples are provided from the L4 dataset, where all cells are pooled from spleen and lung samples. For the LC3 dataset, models were generated for 16 different cell types; 10 from the tumor microenvironment and 6 from healthy tissue. E. Investigation of structural variation across and within different model groups. F. The effect of different RNA-Seq normalization strategies on model similarity across and within model groups.

The presence of misclassified cells is a common problem in single-cell RNA-Seq data, especially when dividing the cells into cell subtype populations, and there are tools available for detecting such cells (11). We investigated to what extent such cells affected the generation of context-specific GEMs by comparing models generated from pure T cell populations to models generated from populations contaminated with varying fractions of cancer cells (Fig 2C). Seemingly, a few percent of misclassified cells have only a negligible effect on model generation compared to other sources of variation (such as data sparsity), while levels of 10-20 % of misclassified cells have a clear negative effect.

A structural comparison of models generated for both bulk and single-cell RNA-Seq profiles from different tissues showed good agreement between models originating from the same tissue and technology (Fig. 2D). However, models generated from similar tissues and different technologies only partly clustered together, suggesting that a combination of technical batch effects and differences in cell type composition between single-cell and bulk have a substantial effect on model generation. Interestingly, immune cells from single-cell lung datasets clustered with GTEx blood samples, which can be expected to have a high immune cell content. We quantified the differences within and across different groups of tissue and technology, which showed that both these variables have a substantial effect on model generation (Fig. 2E).

For practical reasons, context-specific models are often generated from bulk data normalized to transcript per million (TPM), since many other normalization methods are designed to operate on gene counts, and hence do not compensate for gene length. For droplet-based single-cell data this is not a problem, as such data do not need to be normalized by gene length (9). We have previously shown that trimmed mean of M values (TMM) (19) can be applied on TPM data by scaling the TPM values to produce pseudo-counts (9), and we therefore investigated the impact of different normalization methods (Fig 2F). While models generated from bulk data clearly become more similar after TMM normalization, we see no such trend in the single-cell data, which may be explained by the single-cell clusters coming from the same dataset and the same patients. The cell types can therefore be assumed to have been amplified together, and methods beyond library size normalization may therefore be of little use and have a negative effect. Models become more similar across technologies for both normalization methods, and TMM may be a good option for such cases, since the samples still group as expected (Fig. S3). Interestingly, the grouping of the bulk data seems to improve with TMM normalization. The previously observed decreased performance of ftINIT, where the lung models in GTEx were more diverse than for the previous version of tINIT (Fig. S1-S2), was remedied by the TMM normalization, suggesting that ftINIT may be more sensitive to normalization at the chosen gene threshold value. While quantile normalization (20) yields models with even greater similarity, it worsens the grouping on tissue (Fig. S4) and is therefore not recommended.

Another source of variation in single-cell RNA-Seq data that has recently received much attention is variation across samples (Note S1). For example, differential expression analysis with single cell data is known to produce false positives if the variation is measured across cells when not accounting for sample origin (21). In such an approach, the variation across samples is not accounted for, and pooling cells per sample to pseudo-bulk samples followed by applying methods such as DESeq2 (22), which was originally designed for bulk data, remedies the problem. The same problem is faced when trying to estimate the uncertainty in context-specific models generated from single-cell data. The variation across samples is high in single-cell data (Fig. S5) and ideally, procedures that estimate uncertainty should take this into account. In practice, datasets seldom have enough cells to generate reliable models per cell type and sample, and in such cases, we recommend our bootstrapping strategy, although it does not fully account for variation across samples.

### Metabolism across neuron subtypes in the mouse cortex

To assess the utility of our method, we generated context-specific models for different neuron subtypes in the mouse primary motor cortex from a deeply sequenced publicly available dataset (23). Analysis with Seurat (24) yielded a good agreement between the cell subtype definition by Booeshaghi et al and the UMAP projection (Fig. 3A). We selected 17 neuron subtypes for further analysis, each with more than 450 cells in the dataset for further analysis, consistent with our recommendation based on the DSAVE total variation scores (Fig. S6-S7). When using only metabolic genes, the UMAP was still able to separate the dataset per cell subtype (Fig 3B), suggesting that the neuron subtypes exhibit distinct metabolic gene expression signatures that vary more across cell subtypes than within cells of the same subtype.

**Fig. 3:**
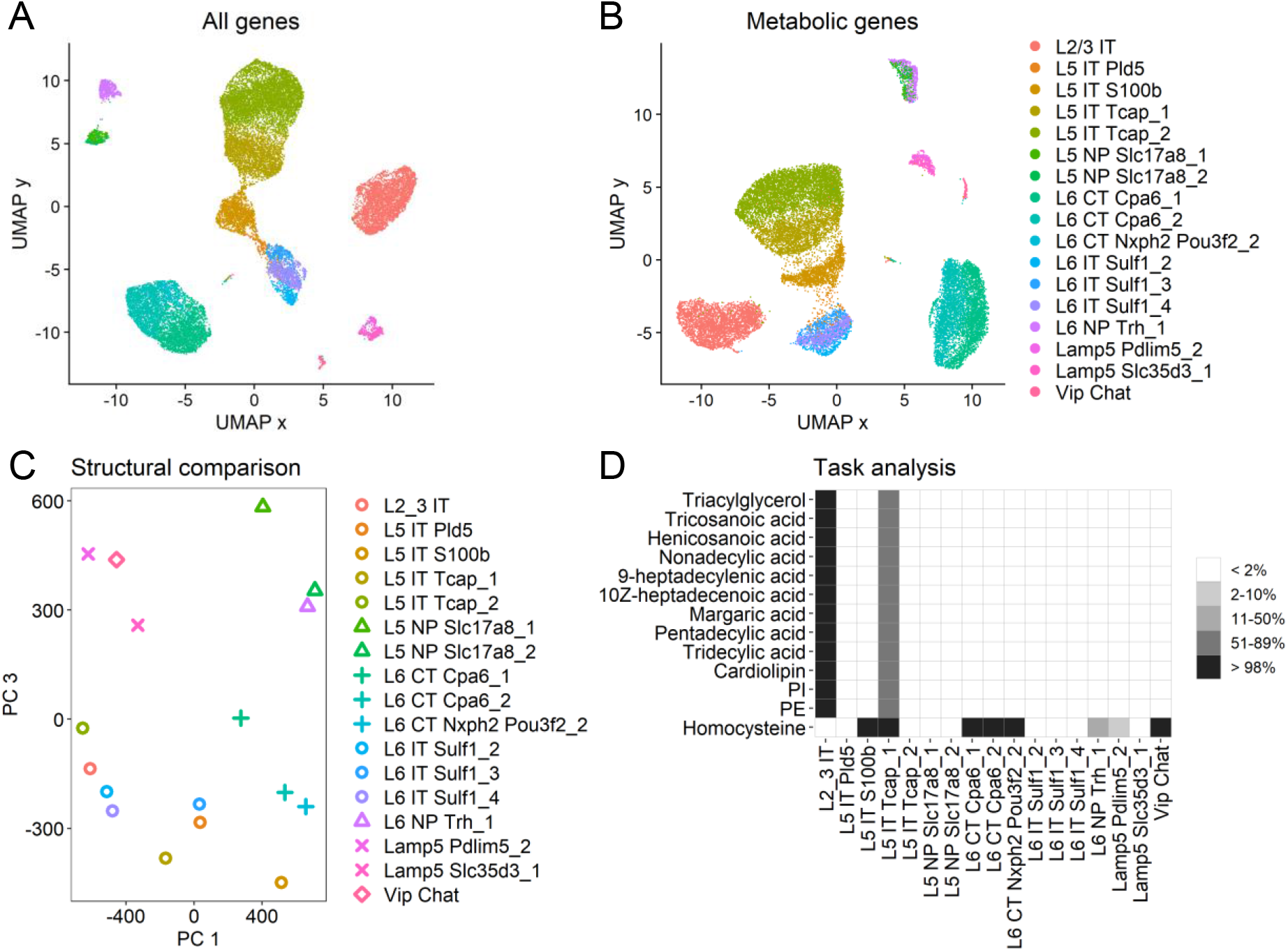
Generation of context-specific GEMs for mouse primary motor cortex cell types. A. Single-cell UMAP projection using all genes, colored by neuron subtype classifications published together with the data. The data displayed is a subset of all cells; only the selected clusters are shown, and only for one batch of the data (with date 4/26/2019). B. Similar to A, but only using the subset of genes present in the Mouse-GEM metabolic model. C. Structural comparison of the context-specific models derived from each neuron subtype. Each reaction is scored based on its presence in 100 bootstrap models, which is used as input to the PCA. PC 2 represent some unknown factor, and is therefore not shown. D. Metabolic task analysis of 100 bootstrap models from each cell subtype. The colors indicate the fraction of the bootstrap models that could perform each task. Only tasks where at least one cell type had more than 98% success rate and at least one had less than 2% such rate are shown. All tasks presented here represents de novo synthesis of the compounds.

To investigate the metabolic networks of the neuron subtypes, we generated 100 bootstrapped single cell populations from each neuron subtype and generated context-specific models for each bootstrap, yielding in total 1,700 models. Since the dataset contains mouse data, we used the Mouse-GEM (25), which is derived from Human1 by gene orthology. The bootstrap models were then pooled together for structural comparison, where each reaction was scored between 0 and 100 representing the number of bootstrap models in which the reaction was present. A PCA analysis revealed structural grouping of neuron subtypes (IT (inferial temporal), NP (near-projecting), CT (corticothalamic), and Lamp5-expressing neurons) when using PC 1 and 3 (Fig. 3B), while we could not see any clear grouping from cortex layer (L2, L5, or L6) (Fig. S8), suggesting that the neuron metabolism is likely defined more by cell function than location. PC2 represents an unknown factor and was therefore excluded from the figure. To quantify the number of reactions that were present in some subtypes but not in others, we defined reactions to be “on” in a subtype if it was present in at least 99 out of 100 bootstrap models, and likewise to be “off” if it was missing in at least 99 out of 100 bootstraps (p < 2.2 * 10^−16^ against the null hypothesis that two reactions, where one is considered “on” and the other “off”, should be equally available; exact Fisher’s test. For statistical considerations regarding multiple testing, see Note S1). A total of 387 reactions (out of 10,376 total reactions) were defined as “on” in at least one cell subtype, and at the same time “off” in at least one other, suggesting a clear distinction between the available reaction networks in the different neuron subtypes. It is also possible to pairwise compare if a reaction statistically has a higher tendency to be “on” in one cell type compared to another, even for reactions that are not considered “on” or “off” (Note S1).

What metabolic capabilities are available to a cell is an interesting property of a cell that can be evaluated by its ability to carry out different metabolic tasks such as *de novo* synthesis or catabolism of important metabolites. We again used our bootstrapped models to perform an analysis of 257 tasks defined in Human1, where we similarly defined “on” if at least 99 out of 100 bootstraps successfully completed the task, and “off” if 99 out of 100 models failed. We found a total of 13 tasks that were considered “on” for at least one cell subtype while “off” for another (Fig. 3C). Most of these differentiating tasks were related to *de novo* synthesis of fatty acids, phospholipids (PI and PE), and cardiolipin. Interestingly, the importance of fatty acids as signaling molecules in neurons has recently been emphasized, and deficiencies in lipid metabolism has been associated with cognitive problems and neurodegenerative diseases (26). The variation of homocysteine synthesis capabilities across neuron subtypes is also interesting. High homocysteine levels in blood are associated with neurological disorders (27,28), and although homocysteine regulation is mainly managed by the liver (27), the ability of some neuron subtypes to synthesize this metabolite suggests that they could play a role in neurological disease. The diversity in homocysteine production capacity among neurons has not been studied, and potential dysregulation of this biosynthetic pathway could therefore be of interest to investigate further.

### Metabolism across cell types in the tumor microenvironment

As a second application, we investigated the diversity in metabolism across cell types in the tumor microenvironment. We downloaded a publicly available lung cancer (lung adenocarcinoma) dataset (29) containing RNA-Seq data from more than 200,000 cells from both healthy lung tissue and tumors originating from 44 patients. The data was first processed using Seurat and the UMAP projections matched well with the cell type classifications provided with the dataset for both cells from the tumor (Fig. 4A) and cells originating from healthy lung tissue (Fig. 4B). The number of UMIs per cell varied substantially across the clusters (Fig. S9A). We estimated the minimum required cluster size to be between 800 -2,000 cells using DSAVE (Fig. S9B) and therefore included the 16 clusters with more than 1,600 cells in the analysis. As expected, the cancer cells showed more diversity than the healthy cell types, since each cancer is unique and has its own transcriptional program (Fig. 4C). Reprocessing the datasets using only metabolic genes yielded similar results, although slightly less separated per cell type, suggesting that each cell type has a unique metabolic program (Fig. S10).

**Fig. 4:**
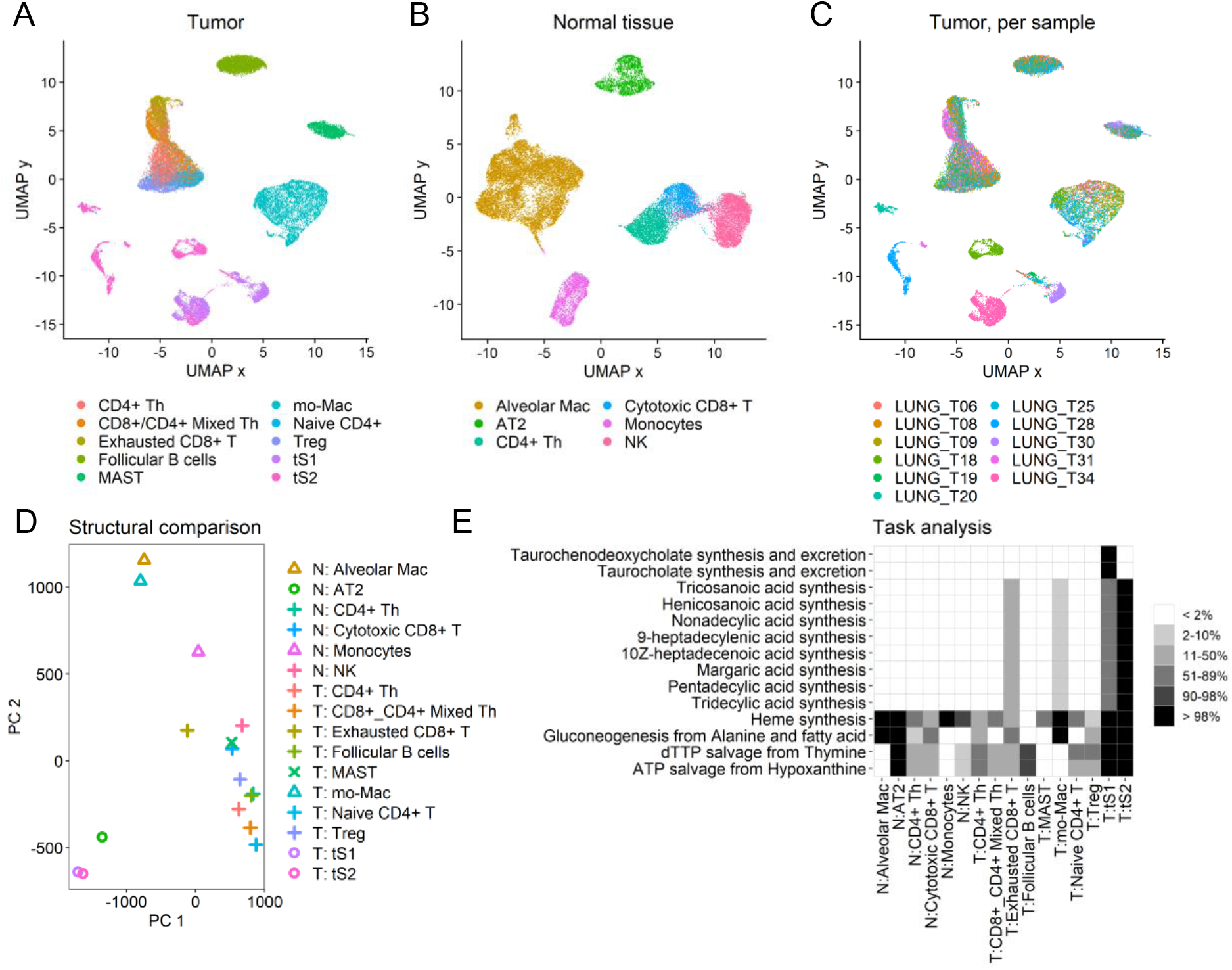
Analysis of the cell types of the tumor microenvironment in lung cancer. A. UMAP projection of cells from tumor tissue. The cells originate from multiple patients. Only cell clusters with at least 1,600 cells are included. The cell type classification used was published together with the dataset. B. Similar to A, but for healthy lung tissue. C. Similar to A, but showing sample origin per cell instead of cell subtype. D. Structural comparison of the context-specific models derived from clusters from both the cancer and healthy tissues. Each reaction was scored based on its presence in 100 bootstrap models, which was used as input to the PCA. The symbol indicates type of cell. E. Metabolic task analysis of 100 bootstrap models from each cluster. The colors indicate the fraction of the bootstrap models that could perform each task. Only tasks where at least one cell type had more than 98% success rate and at least one had less than 2% such rate are shown.

The diversity in metabolism across cell types was first investigated by a structural comparison (Fig. 4D). The cell types roughly clustered into a few groups: myeloid cells (macrophages and monocytes), lymphocytes (T, NK, and B cells) and mast cells, and epithelial cells (alveolar cells and cancer cells), while we could not observe that cell types grouped by tissue of origin (tumor/healthy tissue). In total, 1104 reactions were identified as “on” in at least one cell type and “off” in at least one other type, yielding a diverse set of metabolic networks.

To investigate the differences in metabolic capabilities between cell types, we performed a task analysis on the bootstrap models from all cell types, resulting in 14 tasks that could be confidently completed for at least one cell type while being absent in another (Fig. 4E). At least some tumor cells (tS2) had the ability to generate several types of fatty acids, which is known to be beneficial for tumor progression (30). Interestingly, the two different transcriptional states tS1 and tS2 of the tumor cells exhibited a distinct difference in bile acid metabolism (taurochenodeoxycholate and taurocholate synthesis and excretion), despite that each state was composed of cells from different patients with substantial transcriptional differences. The importance of bile acid metabolism is a topic of recent investigation (31), but its role in lung cancer is not clear and may be of interest for further research. The capacity of the cancer cells to synthesize heme is another interesting observation. The heme synthesis pathway, followed by heme degradation and secretion of bilirubin, provides means to dispose of succinyl-CoA from mitochondria at net production of mitochondrial NADH (32,33). In cell lines with dysfunctional fumarate hydratase, this pathway can be used keep part of the TCA cycle running to generate NADH for OXPHOS, and thereby increase ATP production. Disruption of the heme synthesis pathway was reported to be lethal to such cells (32).

## Discussion

In this work, we developed methods for generating reliable context-specific models from cell populations of single-cell RNA-Seq data. Specifically, we developed a method to estimate the required number of cells per population, a bootstrapping strategy to assess modeling results statistically, and a substantially faster version of tINIT to facilitate the bootstrapping strategy. In addition, we evaluated the effect of normalization methods for the RNA-Seq data and differences in models generated from single-cell and bulk RNA-Seq data. We found that metabolism differs substantially across cell types and subtypes, motivating our approach, and supporting that our methods were useful for finding differences across cell types and could identify metabolic properties known to be associated with the phenotype of interest.

There are many possible ways to investigate metabolism from single-cell RNA-Seq data using genome-scale metabolic models. Reporter metabolites (34) is one method, whereby gene sets are defined based on which metabolites participate in the encoded reaction(s) and used in gene set analysis (GSA). The input to the GSA can for example be p values obtained from differential expression analysis between clusters of single cells as input. Another approach is to penalize reactions based on gene expression, and for different cell populations estimate the total penalty for carrying flux through the reaction network, which is implemented in the COMPASS method (24). The gene expression is in COMPASS estimated per cell by integration over nearby cells, which makes it possible to detect a metabolic switch in either a cell continuum or between clusters. While these methods were proven useful (24,25,35), they are designed to directly detect upregulated pathways, while our method generates a model that can be used for further simulations, which enables the investigation of other questions. In this study, we showcased our method using metabolic task analysis, but it also allows for more advanced modeling approaches. Such methods could involve the use of metabolite uptake constraints (e.g., based on diffusion), constraints on enzyme usage, or simulations involving the interplay between several cell types (4,36).

Statistical inference is often a challenge when using single-cell RNA-Seq data – single-cell datasets often do not contain enough samples, or enough cells per sample, to apply statistics in a similar way as for bulk RNA-Seq samples. While our method partly suffers from the same weakness, our bootstrapping approach provides some statistical assurance, although subject to certain assumptions (Note S1). It is important to note that applying our method over cells from a mix of several samples requires that the cell type proportions are reasonably similar across samples, since batch effects between patients could otherwise bias the results, and a prefiltering of cells to ensure cell type proportions may be necessary.

Although some methods have recently emerged (15,24,34), the use of GEMs together with single-cell RNA-Seq data to study disease is still in its infancy. However, such analyses hold great potential – single-cell RNA-Seq enables characterization of all cell types in the human body (37). In addition, each cell type can come in various states, and sometimes continuums, and such aspects are difficult to capture in bulk data, even for FACS-sorted cell populations. We have shown that the metabolic transcriptional program varies substantially across cell types, suggesting that study of individual cell types will provide further detail when studying metabolism in complex organs. With further development of both single-cell RNA-Seq and GEMs, the combination of the two holds promise for a substantial contribution in unraveling the key metabolic features in human health and disease.

## Methods

### Datasets

To evaluate our methods, we downloaded 8 different single-cell RNA-Seq datasets (29,37–43). Bulk RNA-Seq data was downloaded from GTEx (18) for comparison of modeling performance across tissues. For the gene essentiality evaluation, we downloaded bulk RNA-Seq TPM data and CRISPR screening data for gene essentiality from DepMap (16,17). To compare the variation between pools of single cells and bulk RNA-Seq, we used the same FACS-sorted T cell samples as previously used for DSAVE, available from the BLUEPRINT epigenome project (11,44). Detailed information about the datasets is available in Table S1.

### Model preparation

To reduce the size of the genome-scale metabolic model, and thereby the execution time of both versions of tINIT, we removed in total 1,496 reactions from Human1 that were deemed unnecessary, leaving a model with 11,582 reactions. The reactions removed included reactions related to drug metabolism, amino acid triplets, and a list of reactions that were identified as duplicates. The latter were also permanently removed from the Human1 model as a curation step for future modeling work.

### ftINIT

The ftINIT method is described in detail in Note S2. In short, ftINIT runs in three steps: 1) Simplified run where secretion of metabolites is allowed and many reactions are omitted from the problem (many reactions without GPRs, reversible reactions with positive scores). 2) Run where the reversible reactions are included, and the reactions turned on in step 1 are excluded. 3) Run where metabolite secretion is no longer allowed and where most reactions without GPRs are included in the problem. Step 3 was omitted for the generation of all models used in this work. In step 1 and 2, the model was simplified by removing the following metabolites: ‘H2O’;’Pi’;’PPi’;’H+’ (except for OXPHOS);’O2’;’CO2’;’Na+’. The ‘rxns to ignore mask’ was set to [1 1 1 1 1 1 1 0], which effectively means that a collection of reactions without gene rules, including spontaneous reactions, exchange reactions, transport reactions, and custom reactions are omitted from the optimization problem and are always included in the final model. The custom reactions were in this study selected to include reactions for protein generation, reactions that pool metabolites, and a few reactions handling radicals (which we interpreted as spontaneous), and a few additional reactions that we strongly suspect are spontaneous, in total 69 reactions. None of the custom reactions had gene rules.

ftINIT is designed to work with the Gurobi solver. ftINIT was run with with a MipGap of 0.04% and a time limit of 2 minutes per step, and has not been tested with other solvers, although it may work. The MipGap parameter describes a limit for the maximal possible error of the objective function as calculated by the solver – if the maximal possible error becomes smaller than this value the solver stops, reporting that a solution is found. The time limit parameter makes the solver stop after a certain time, regardless of the status of the solution. Should a solution not be found within the time limit of 2 minutes, the allowed MipGap is increased to 0.30%. If the MipGap after two minutes is higher than the allowed value, the optimization will be rerun with a time limit of 5,000 seconds – though based on our observations this only happens on rare occasions. The parameters for the previous tINIT could be freely specified but was often used with only a time limit parameter of either 1,000 s, 5,000 s, or 10,000 s depending on the complexity of the samples.

### Pooling single-cell samples to simulate pseudo-bulk

The single-cell data was pooled into pseudo-bulk RNA-Seq profiles for use with ftINIT/tINIT. We chose a strategy that assigned each mRNA molecule equal importance, and therefore summed all UMIs from all cells in a population into a single pseudo-bulk sample. The sample counts were then scaled to a total sum of 10^6^ (counts per million, CPM, normalization), except in cases where other normalization methods such as TMM or quantile normalization were applied.

### Evaluation of tINIT execution time

tINIT and ftINIT were run for the 10 first tissues in the GTEx dataset, where each tissue was represented by the median expression values of all RNA-Seq samples from that tissue. The time was measured per run, performed on a standard laptop computer (Intel Core i7-6600U, 2.60 GHz, 2+2 cores). ftINIT uses a MipGap of 0.04% for the first two minutes of each step, which is increased to 0.30% after two minutes if the solver has not found a solution within that time. Thus, since the worst-case scenario of ftINIT can lead to an acceptance of a MipGap of 0.30%, we set the optimization for tINIT to end at a MipGap of 0.30%. This is not the set of parameters by which tINIT is normally run (where only a time parameter is set), but this set of parameters makes the execution times comparable between the methods, although in favor of the old version.

### Evaluation of required pool size and cell type contamination

For ftINIT, a total of 10 different single-cell pool sizes were examined. For each pool size and dataset, we generated 20 pairs of random cell populations of the examined pool size, where the two populations in each pair do not share any cells. The cells in the populations were pooled to generate RNA-Seq profiles and processed using ftINIT, generating 400 models per dataset. Each pair was then compared for reaction content using Jaccard similarity coefficient. The mean value of the Jaccard similarity coefficients was then calculated for all pairs within the same pool size and dataset.

For the cell type contamination investigation, a similar approach was taken as for the ftINIT investigation. We selected two cell populations from patient 4 of the LC dataset: one containing T cells (LC T) and the other containing malignant cells (LC M). 8 different pool sizes were determined and in addition 5 different fractions of contamination. 60 pairs of random cell populations were then generated for each pool size and dataset. The first of the cell populations in each pair was uncontaminated, drawn completely from the LC T cell population. The second population was drawn from both LC T and LC M, such that

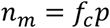

where *n*_*m*_ is the number of cells drawn from the LC M population, *f*_*c*_ is the fraction of contaminated cells to include, and *p* is the pool size. The remaining cells up to *p* cells were drawn from the LC T population. Instead of running ftINIT (to reduce computation time) we generated reaction scores (see Note S2) for each reaction in both populations of the pair, and estimated each reaction to be “on” if the reaction score was larger than 0, and otherwise “off”. The Jaccard similarity coefficient was then estimated for each pair. The mean values for the Jaccard similarity coefficients were then estimated per pool size and fraction of contamination.

### t-SNE and PCA (when used separately, not with Seurat)

The tsne function in MATLAB was used to generate t-distributed stochastic neighbor embedding (t-SNE) (45) coordinates from the model structure. PCA was performed using the function “prcomp” in

R. The t-SNE figures were based on the presence of each reaction in a single model, and the Hamming distance was used as distance between samples. For the PCA plots, each reaction was scored between 0 and 100 depending on how many bootstrap models included the reaction. The PCA was then performed with these scores per reaction as variables across the cell subtypes.

### Calculation of Jaccard index for reaction scores

To avoid the generation of large quantities of context-specific GEMs in Fig. 2C, we as an alternative to investigating the structural difference between pairs of context-specific models instead generated reaction scores for each gene in each pooled RNA-Seq sample. It is then assumed that there for each gene is one reaction that depends on that gene only. The reaction score *r*_*g*_ for each gene *g* is calculated as

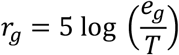

where *e*_*g*_ is the gene expression in CPM for gene *g* and *T* is the threshold value, which is set to 1. To calculate the Jaccard index, each gene *g* is treated as “off” if rg < 0 and “on” otherwise. This scoring is very similar to that performed in ftINIT. The Jaccard index is then calculated on such vectors with “on” (1) and “off” (0) values between pairs of samples.

### Single-cell analysis using Seurat

Seurat v. 4.04 (24) was used to analyze the single-cell datasets MCOR3 and LC3, using a standard Seurat pipeline, including log normalization (scale.factor=10,000), finding of variable genes(vst, 2,000 features), data scaling, PCA, finding of neighbors (using 15 PCs), clustering (resolution = 0.5, otherwise standard parameters), followed by UMAP (using 15 PCs) and DimPlot for visualization. The cell type classifications provided as part of the publications were used, and all cells that had a classification matching one of the cell types/subtypes selected for study were included in the processing. No other cell filtering was performed. For the LC3 dataset the healthy tissue data and tumor data were processed separately. The metabolic genes for human and mouse were extracted from the respective models, where all genes included in a gene rule was considered metabolic. Genes from 5 reactions were excluded (‘MAR09577’, ‘MAR09578’, ‘MAR09579’, ‘MAR07617’, and ‘MAR07618’) since these reactions describe phosphorylation of proteins, which we consider to be signaling reactions. In total, we classified 2,784 genes in MCOR3 dataset and 2,912 in the LC3 dataset as metabolic.

## Software

The data was analyzed using MATLAB R2019b and R version 4.1.1. To ensure the quality of our analyses, we verified and validated the code using a combination of test cases, reasoning around expected outcome of a function, and code review. The details of this activity are available in the verification matrix available with the code.

## Declarations

## Availability of data and materials

The model Human1 is available in GitHub (https://github.com/SysBioChalmers/Human-GEM). The processed data and source code are available in Zenodo: https://doi.org/10.5281/zenodo.6482794. The source code is also available in GitHub: (https://github.com/SysBioChalmers/SingleCellModeling). The ftINIT method is planned to be implemented in RAVEN Toolbox (46).

## Authors’ contribution

J.G, J.R, and J.N. planned the project. J.G. wrote all software and made all analyses and figures. J.G. wrote the draft manuscript. All authors reviewed and edited the manuscript. J.R. and J.N. supervised the project. J.N. acquired funding for the project.

## Acknowledgements

The computations were performed on resources provided by the Swedish National Infrastructure for Computing at C3SE.

