## Supplementary information for "Generation and analysis of context-specific genome-scale metabolic models derived from single-cell RNA-Seq data"

### Supplementary Figures

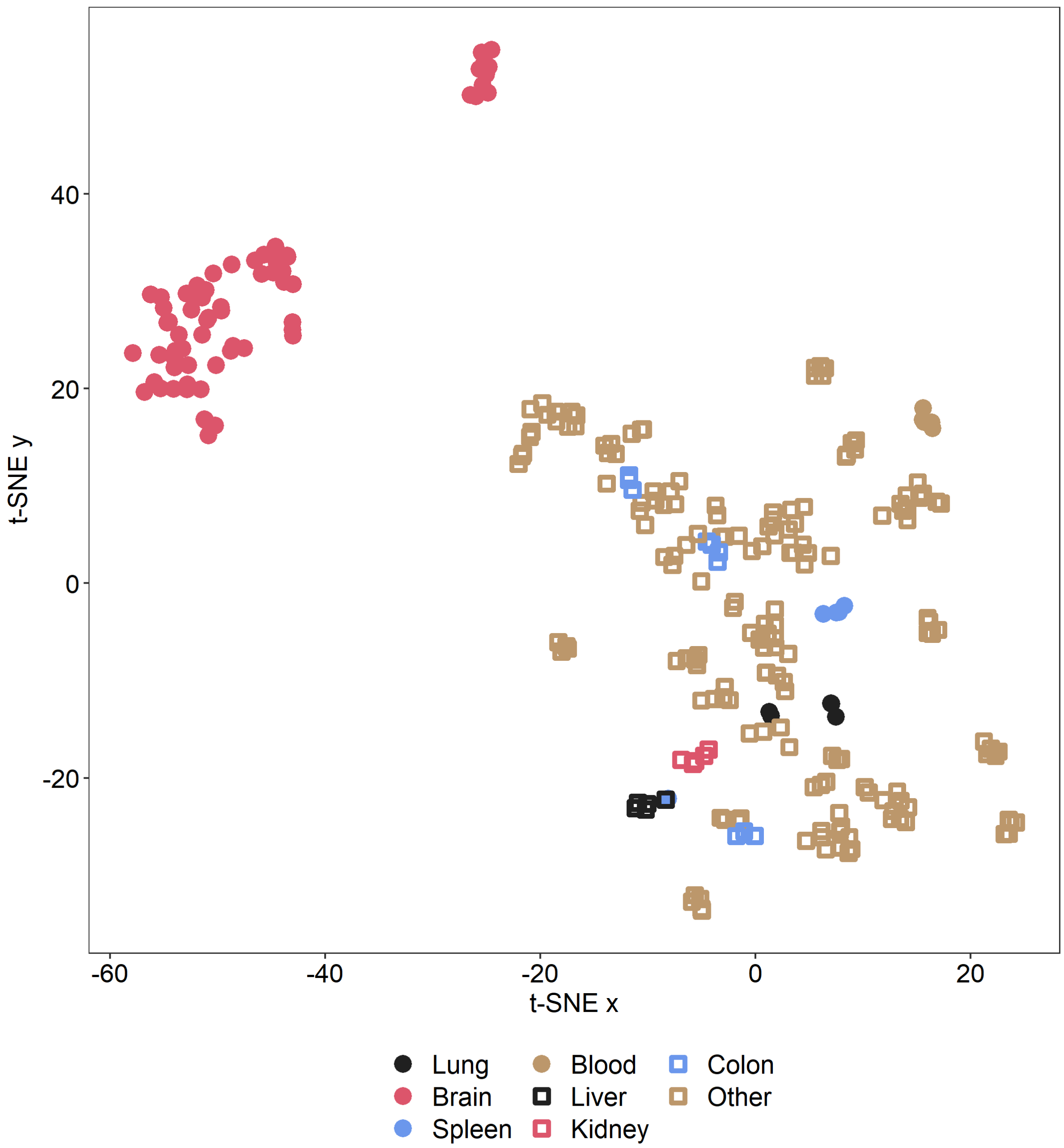


***Fig S1: Grouping per tissue for the previous version of tINIT.*** *Structural comparison of genome-scale metabolic models generated by tINIT from RNA-Seq profiles from GTEx (bulk data), 5 samples per tissue, displayed as a t-SNE projection.*

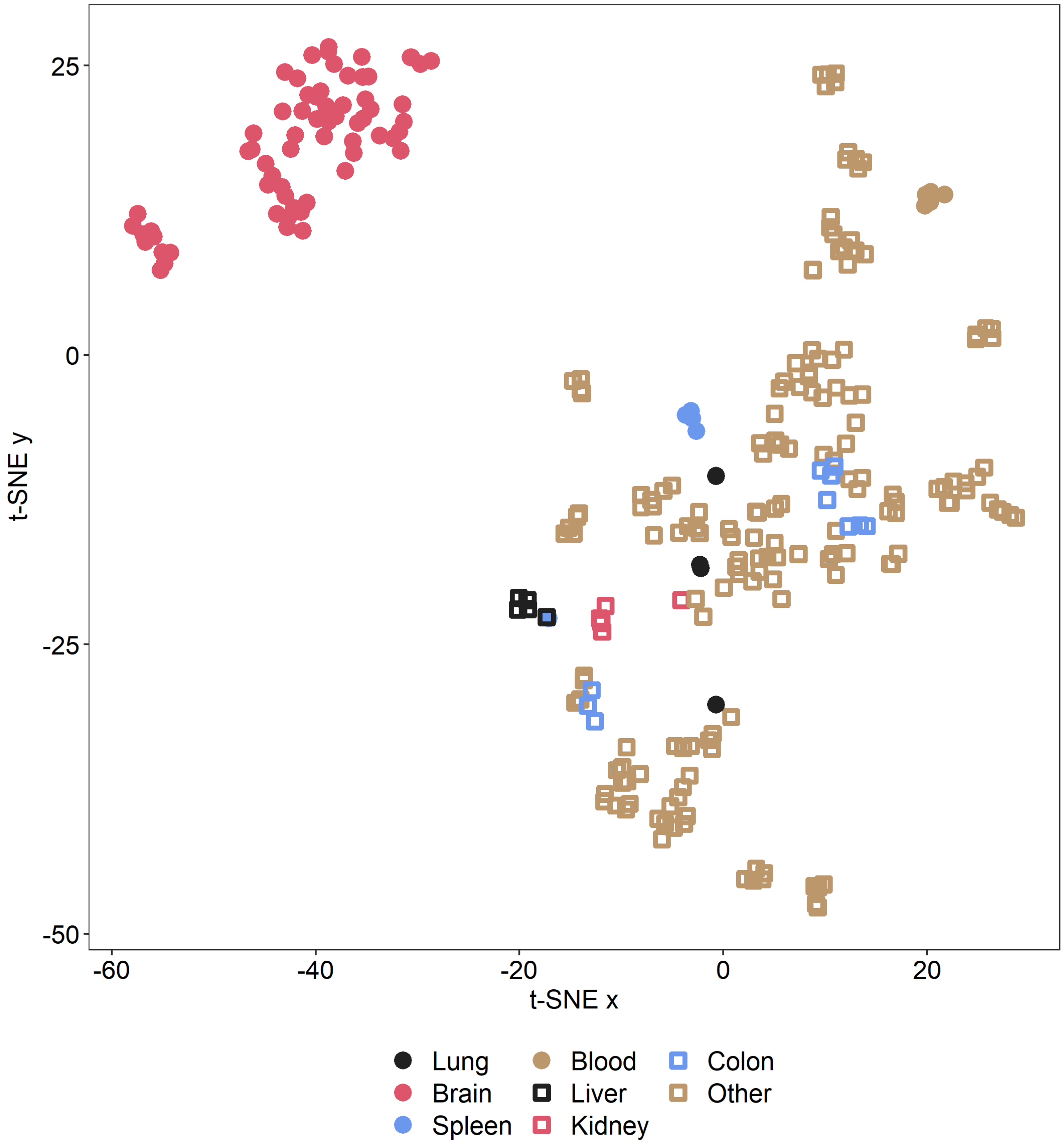


***Fig S2: Grouping per tissue for ftINIT, the new faster version of tINIT.*** *Structural comparison of genome-scale metabolic models generated by ftINIT from RNA-Seq profiles from GTEx (bulk data), 5 samples per tissue, displayed as a t-SNE projection. The grouping is very similar to that of the old tINIT, but with slightly more spread for kidney and lung samples.*

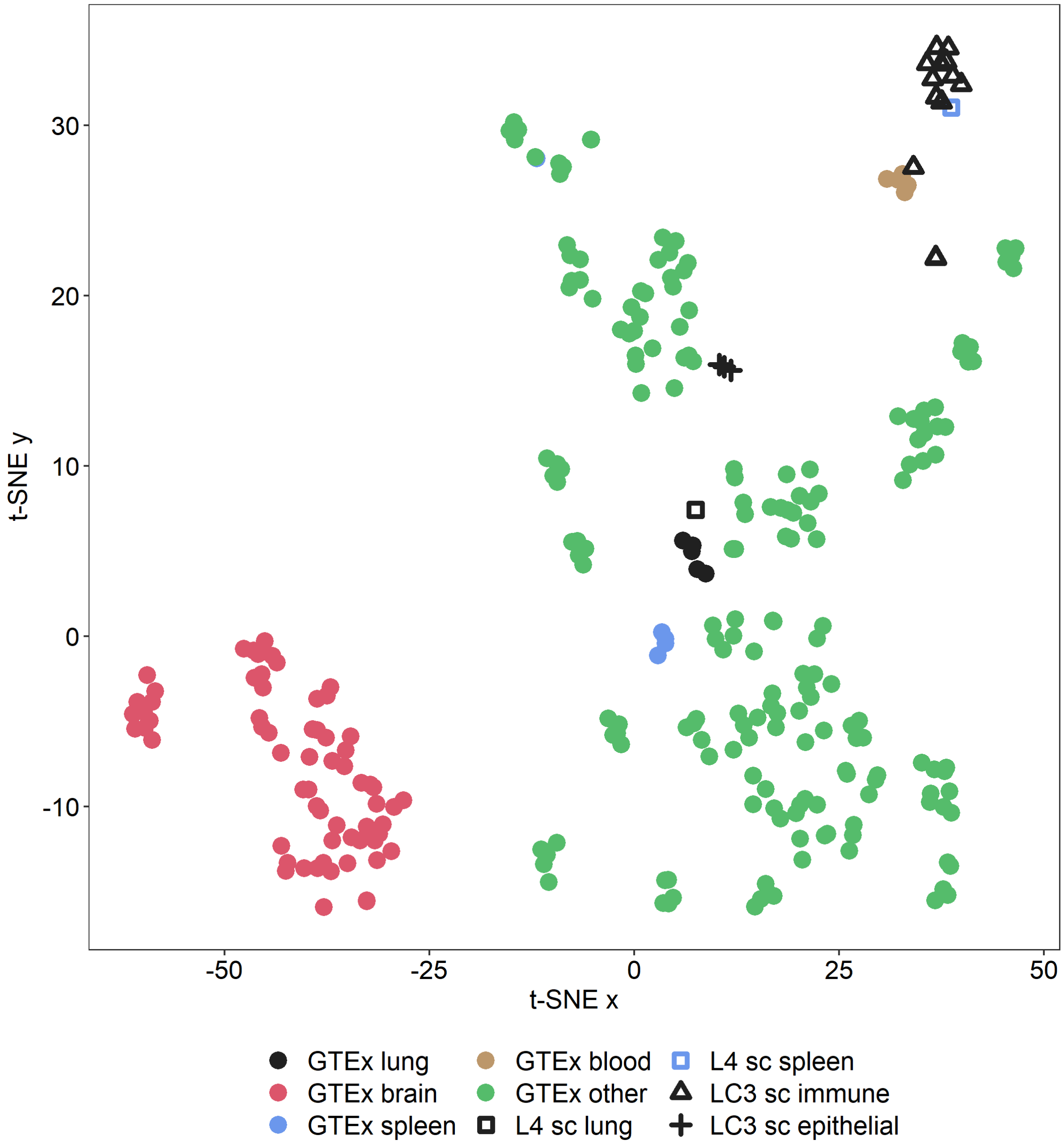


***Fig. S3: Effect of TMM normalization on structural comparison.*** *Structural comparison of genome-scale metabolic models generated by ftINIT from RNA-Seq profiles from GTEx (bulk data) and several single-cell RNA-Seq datasets, displayed as a t-SNE projection. See Fig. 2D in the main text for details about the datasets used.*

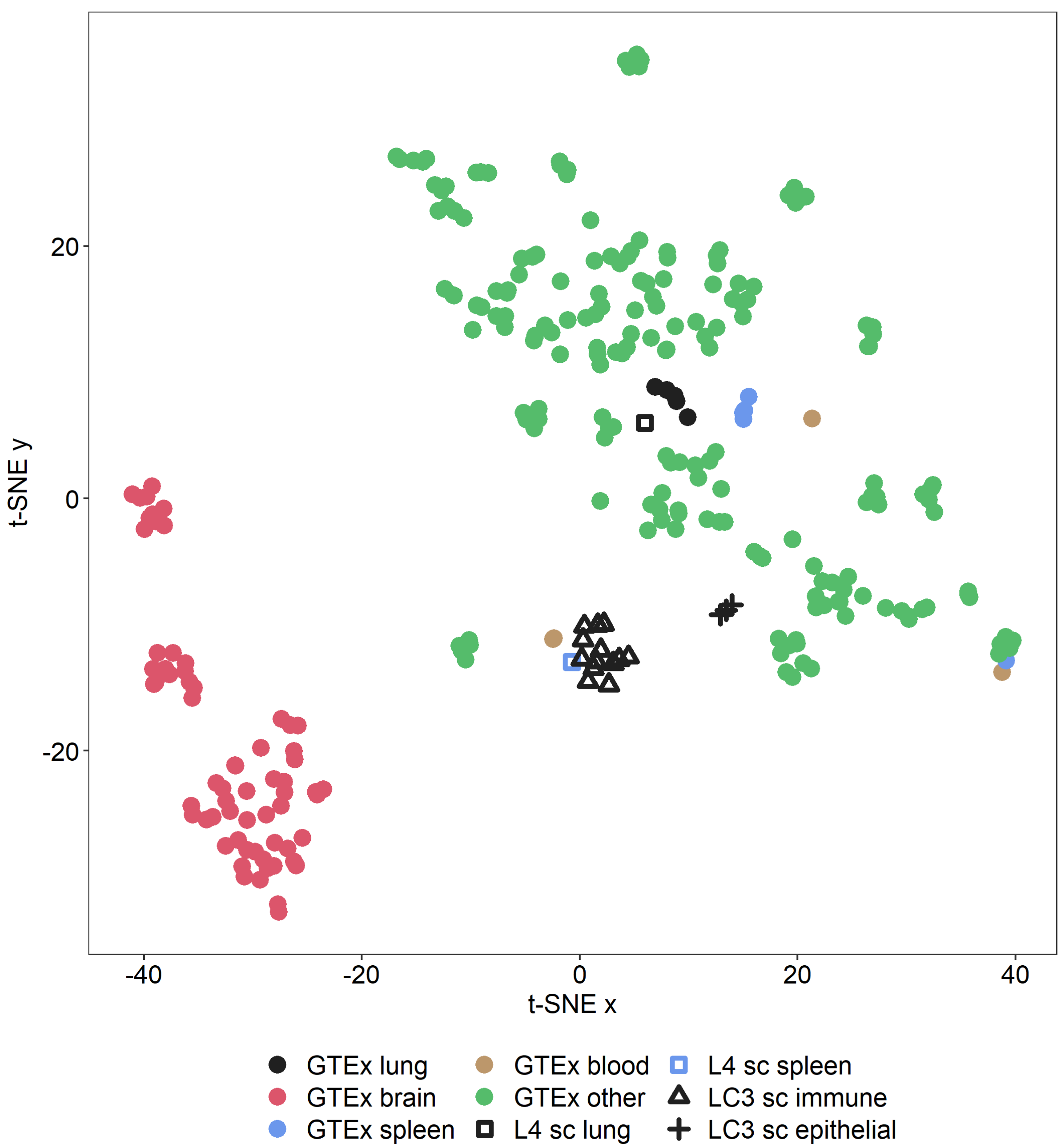


***Fig. S4: Effect of quantile normalization on structural comparison.*** *Structural comparison of genome-scale metabolic models generated by ftINIT from RNA-Seq profiles from GTEx (bulk data) and several single-cell RNA-Seq datasets, displayed as a t-SNE projection. The GTEx blood samples are no longer grouped. See Fig. 2D in the main text for details about the datasets used.*

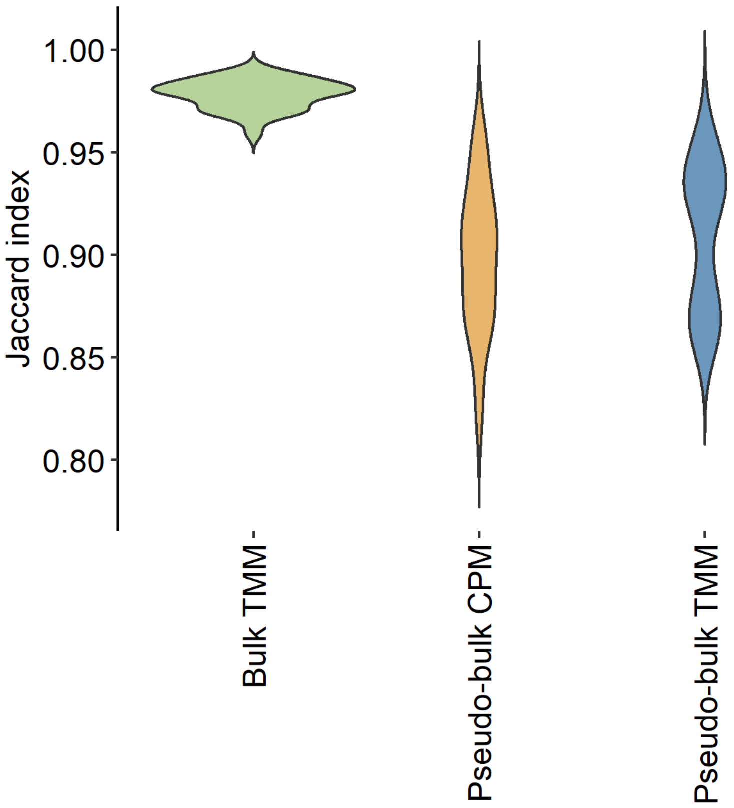


***Fig. S5: Variation across samples in single-cell data.*** *The Jaccard index is calculated between all pairs of 8 samples of each category (bulk T cells, TMM normalized, the Bulk T cell dataset; single-cells pooled per sample, T cells from the HCA CB dataset, CPM normalized; single-cells pooled per sample, T cells from the HCA CB dataset, TMM normalized).*

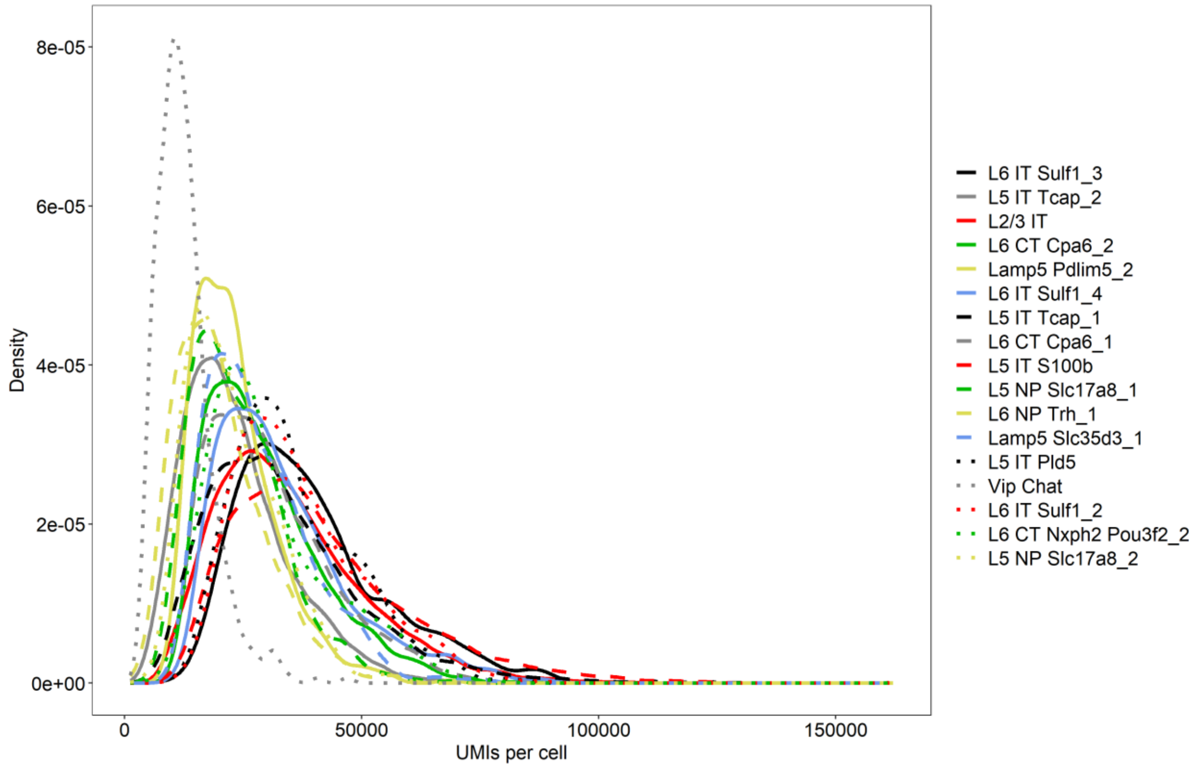


***Fig. S6: Distribution of UMIs per cell per cluster in the MCOR3 dataset.***

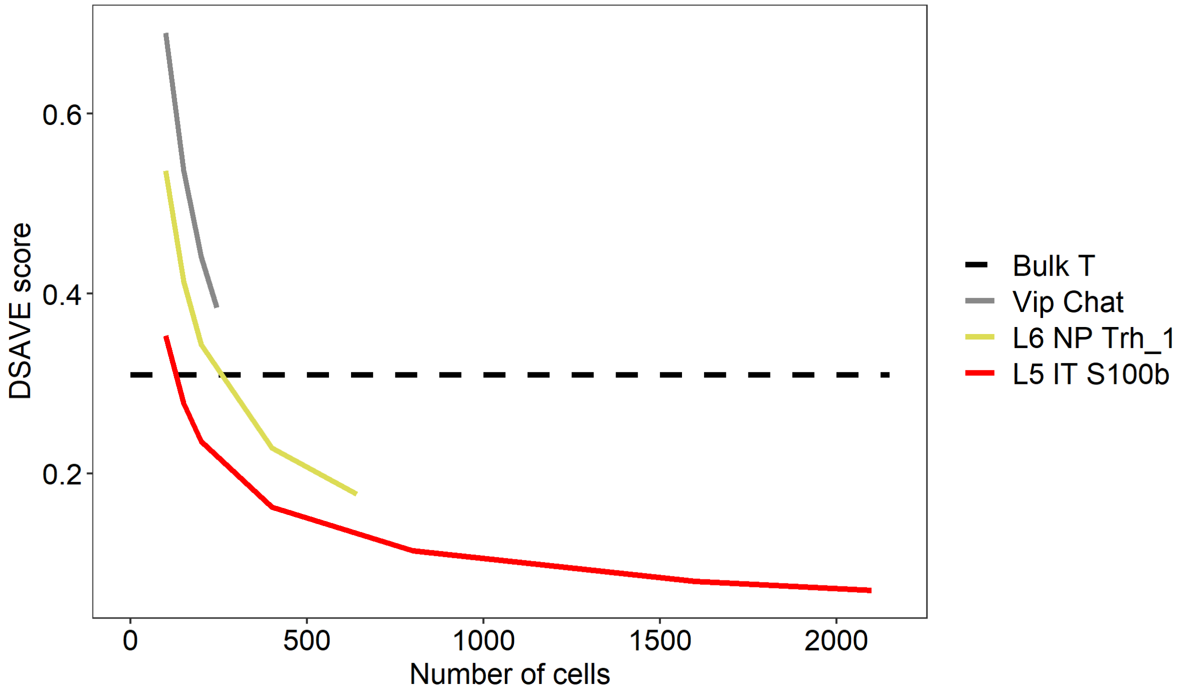


***Fig. S7: DSAVE analysis of the MCOR3 dataset.*** *Only a subset of the clusters is shown for clarity; the Vip Chat is the cluster with the least UMIs per cell (see Fig. S6). Although we do not have enough cells for prediction when this cluster reaches the bulk variation, since DSAVE requires 2 times the number of cells to be able to measure the variation, we estimate it to be less than 450 cells.*

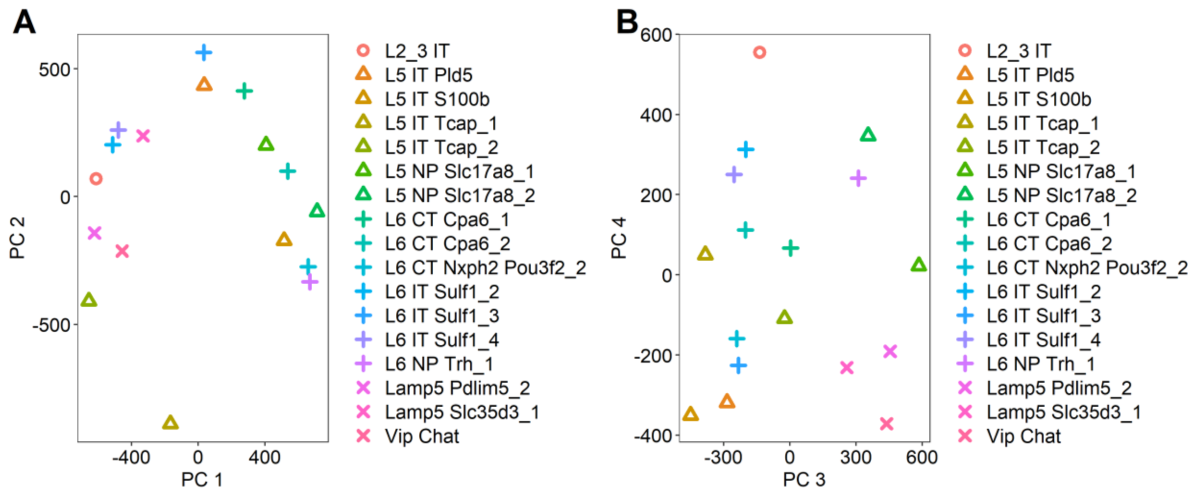


***Fig. S8: Neuron metabolic network dependency on cortex layer.*** *Structural **c**omparison of context-specific models generated by ftINIT from pooled single-cell clusters from the MCOR3 dataset, visualized as a PCA projection. A. The two first principal components. No clear dependence on cortex layer. B. PC 3 and 4, no clear dependence on cortex layer.*

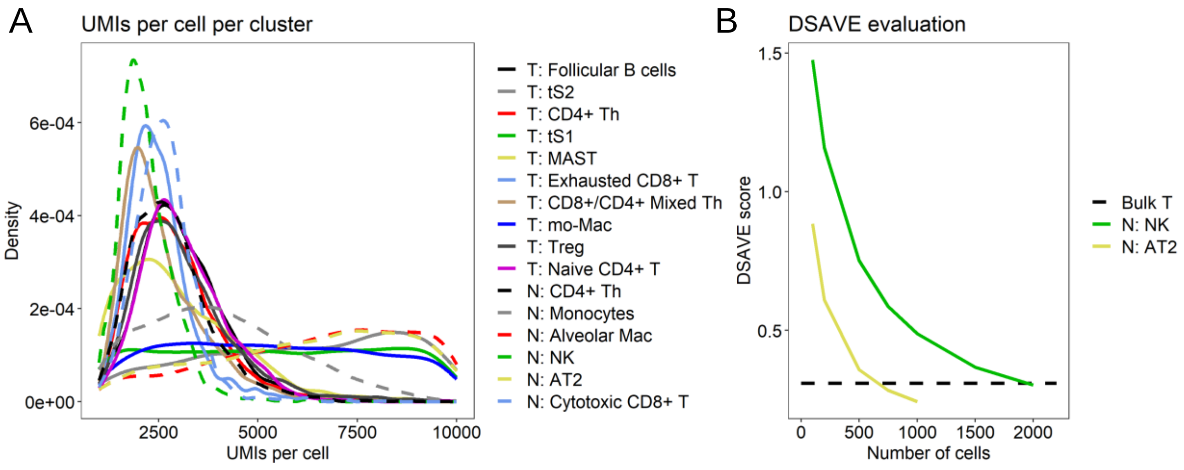


***Fig S9: Evaluation of cluster size for the LC3 dataset.*** *A. The distribution of UMIs per cell across clusters. B. DSAVE total variation score for two clusters with high (N: AT2) and low (N: NK) average number of UMIs per cell.*

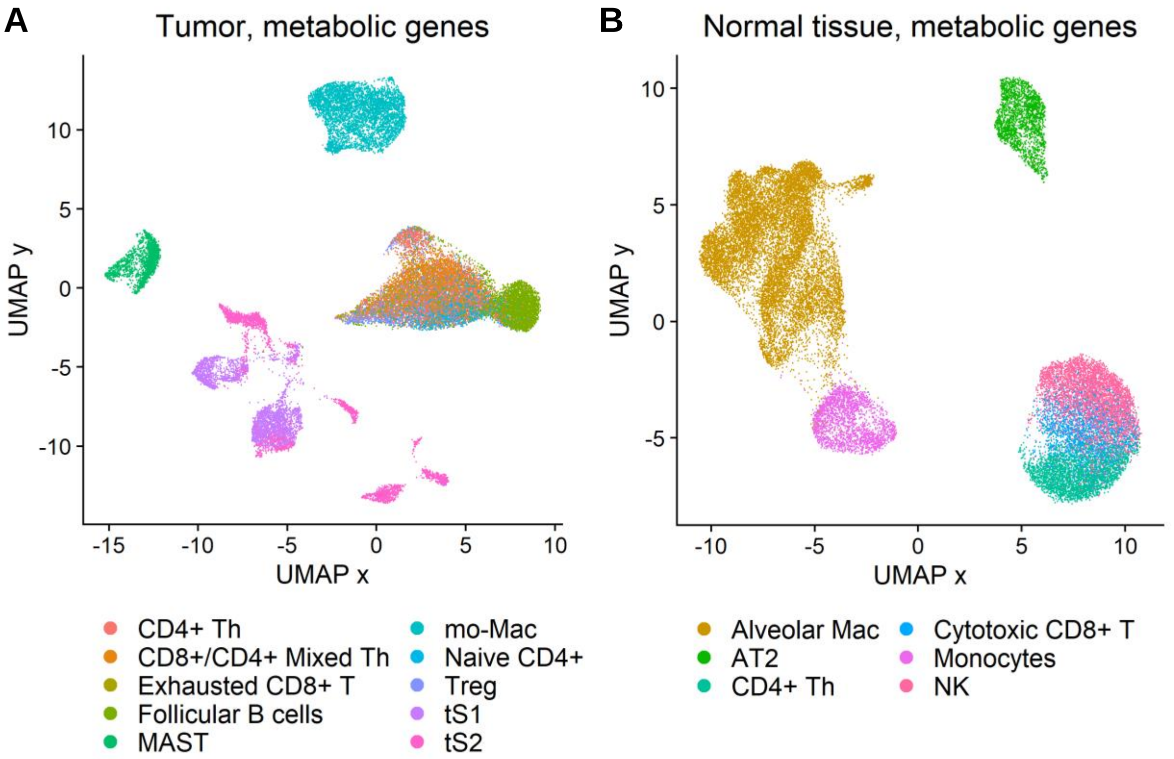


***Fig S10: UMAP projection using only metabolic genes for the LC3 dataset.*** *These figures are similar to Fig. 4 A-B in the main text, with the difference that only metabolic genes were included in the data processing. A. Tumor samples. B. Samples from normal tissue.*

### Supplementary Tables

***Table S1. Datasets***

| ID | Information |
|---|---|
| HCA CB | Cord blood data from Census of Immune cells, Human Cell Atlas (1,2), in total ~254,000 cells from 8 patients, 10X Chromium v2. The data can be downloaded from Census of immune cells. https://data.humancellatlas.org/ The T and B cells of this dataset, identified using the cell type classification of the authors, are referred to as HCA CB T and HCA CB B, respectively. For practical reasons, these populations have been reduced to 25,000 cells each. |
| PBMC68k | ~68,000 blood (PBMC) cells from a single patient, 10X Chromium v1, denoted as Fresh 68k PBMC (3). The T cells of this dataset, as identified by the authors, are referred to as PBMC68k T. The data is available at 10X Genomics home page (https://www.10xgenomics.com/resources/datasets/fresh-68-k-pbm-cs-donor-a-1-standard-1-1-0). |
| Mel | 4,600 cells from the tumors of 19 melanoma patients, Smart-Seq2 (4). The T cells of this dataset, as identified by the authors, are referred to as Mel T. The data is available for download at the GEO data repository, accession number 72056. |
| LC | ~39,000 cells from the tumor microenvironment of lung cancers and ~13,000 cells from adjacent healthy lung tissue, 10X Chromium (mix of v1 and v2) (5). The T cells and macrophages, as identified by the authors, are referred to as LC T and LC Mac, respectively. The data is available for download in ArrayExpress under accessions E-MTAB-6149 and E-MTAB-6653. |
| TCD8 | ~10,000 CD8 positive FACS-sorted T cells from the blood (PBMC) of a single patient, 10X Chromium v2 (6). For symmetry, the cells in this dataset are referred to as TCD8 T. The data is available for download at the GEO data repository, accession number 112845. |
| LC3 | ~100,000 cells from lung tumors and ~100,000 cells from adjacent healthy lung tissue, in total 44 patients, 10X Chromium v2 (7). The data is available for download at the GEO data repository, accession code GSE131907. |
| L4 | ~57,000 cells from lung tissue and ~94,000 cells from spleen tissue, 10X Chromium v2 (8). All cells from the lung and spleen sample, respectively, were pooled to form two pseudo-bulk samples. The data is available through the Human Cell Atlas Data Coordination Platform and NCBI BIOPROJECT accession code PRJEB31843. |
| MCOR3 | ~94,000 deeply sequenced cells from the mouse primary motor cortex, primarily neurons, 10X Chromium v3 (9). The data is available at http://data.nemoarchive.org/biccn/lab/zeng/transcriptome/scell/ |
| Bulk T | 8 T cell bulk RNA-Seq samples from human PBMC, available from the BLUEPRINT Epigenome Project(10). The samples were taken from the project EGAD00001001173 and have the following sample IDs: S002EV11, S004M711, S007DD11, S007G711, S008H111, S009W411, S001FRB1, and S0041C11. The FASTQ files |

| | |
|---|---|
| | were processed as previously described (11): The FASTQ files were first processed using kallisto (12) to produce gene counts (estimated counts produced by kallisto). The data was then normalized using TMM (13), and scaled to an average count of $10^6$ per sample. |
| DepMap | Bulk RNA-Seq TPM-normalized data and CRISPR screening data for gene essentiality from DepMap (14,15) (version 21Q3, specifically "CCLE_expression_full.csv" and "Achilles_gene_effect.csv"). Only 15 samples in the RNA-Seq data with matching gene essentiality data were used. The data is available from https://depmap.org/portal/. |
| GTEx | Bulk RNA-Seq data from in total 53 tissues from GTEx (16) (version 8, RNA-Seq v. 1.1.9, both TPM and raw counts files). Only 5 samples from each tissue were used. For the execution time evaluation, the median expression within each tissue was used. The data is available from the GTEx portal, https://gtexportal.org/home/. |

.

### Supplementary notes

#### Supplementary note 1 – Statistical considerations for generation of context-specific models from single-cell RNA-Seq data

A central question when generating context-specific models from single-cell RNA-Seq data is how to assess the uncertainty both in the data and in the algorithm. What we primarily seek to estimate when generating context-specific models from single-cell data is the uncertainty of the presence of reactions (or other binary aspects, such as the ability to perform metabolic tasks). Given several cell populations per cell type, it is possible to do a pairwise comparison between two cell types, where a statistical test could explain if the tendency to include a reaction is higher in models generated from one cell type than the other. However, for cases where we seek to compare multiple samples, it would also be beneficial to estimate the uncertainty in reaction presence in a sample without comparing it to other samples. In this note, we explain how we address these two cases statistically, especially the latter.

In the ideal case, an adequate number of biological samples with enough cells per cell type is available in the dataset. Differences in reaction presence between two samples can then be tested on pooled pseudo-bulk samples using a two-sided Fisher's exact test, followed by correction for multiple testing by for example the Benjamini-Hochberg method. However, most datasets do not contain enough cells and samples to obtain stable models and significance using such a method, which poses a challenge for the statistical analysis.

The variation between two pools of cells originating from different samples can be divided into two components: 1) systematic variation (technical and biological) between the biological samples and 2) cell-to-cell variation (mainly sampling effects but also effects such as transcriptional bursting), which we assume is mostly independent of sample. The first component is independent of pool size, while pool size heavily influences the second. At an infinite pool size, the second component is zero, whereas it dominates the variation for small pool sizes. The importance of the two components also varies with gene expression; for highly expressed genes, the second component is less important than for lowly expressed genes. As an example of the importance of these components, Squair et al found that differential expression analysis methods that operate on pseudo-bulk pools of single cells per sample are more accurate compared to methods operating directly on single cells, ignoring sample origin (17). They show the main source of the error to be the inability to account for systematic variation across samples in the cell-wise methods, leading to an underestimation of the total variation, giving rise to false positives. The effect was only detected for highly expressed genes, which is likely due to the greater contribution of sampling effects (component 2) to the total variation compared to systematic across-sample variation (component 1) for lowly expressed genes, and therefore the total variation is only slightly underestimated by mixing samples for such genes.

For datasets not containing enough cells and samples to enable statistical tests across pseudo-bulk samples, we propose to treat all cells belonging to a certain cell type as a single population, regardless of sample origin. As an approximation, we assume that the available cells in the population are a perfect representation of the cell type, containing all possible cell variants in proportions representative of the cell type. We can then create bootstraps from the cell population to estimate the variation in the cell population and pool each bootstrap to generate a pseudo-bulk sample. This approach makes it possible to statistically assess the presence of reactions in smaller datasets and is a good option in cases where insufficient data is available for tests across samples. However, this approach comes with two drawbacks: 1) the variation is underestimated since component 1 of the variation is not included, which could lead to false positives; 2) the variation is underestimated since the available cells in the population is not a perfect representation of the cell type, which also could lead to false positives. The latter effect worsens with small pool sizes. As an example, a pool of 10 cells will have many genes that are falsely not expressed due to sampling effects. Bootstrapping will not yield any variation in gene expression for such genes, and associated reactions will confidently, but falsely, be considered non-present. So, while the bootstrapping method is useful, it is important to understand that it will produce more false positives than a test across samples and that the pool size has an effect. If the pool size is small, many genes will falsely be zero (or very high), while if it is large, the variation considered to originate from component 2 may be considerably smaller than that of component 1, which is largely ignored. It is also worth noting that in the case where uncertainty is estimated across samples, differences in the number of cells (and counts) between cell types could lead to biases in a similar way to drawback 2 explained above. An alternative explanation is that the uncertainty in the estimated variables (on/off) cannot be estimated directly from the data, in contrast to the case of bulk RNA-Seq differential expression analysis, where the uncertainty can be directly estimated from the counts assuming they follow a negative binomial distribution for each gene.

We seek to be confident in that a reaction considered present in one cell type statistically shows a larger tendency to be present in that cell type compared to cell types where it is considered non-present. Since we test all reactions, the result needs to be corrected for multiple testing. In this work, we used the bootstrapping strategy explained above with 100 bootstraps, and therefore seek to statistically compare the number of bootstraps for which the reaction is on between two cell types. Ideally, we would like to find a number *x* of bootstraps such that if a reaction is present in x bootstraps in one cell type, the reaction should with statistical significance be more available in that cell type compared to another cell type where the reaction is present in 100-*x* bootstraps. Finding such a number would enable us to define reactions as “on” and “off” in cell types and move away from relative pairwise comparison of cell types, where we are just be able to determine if a reaction is more present in one cell type than another. However, when using the Benjamini-Hochberg procedure to adjust p values for multiple testing, the results of such an adjustment will depend on the p values from other reactions for the pair of cell types compared, and the value of *x* would be different for different pairs of cell types. It is thus difficult with this definition to determine if a reaction is present with a significant p value without relating it to another cell type, which would be desired. We have therefore adopted the strategy to define fixed thresholds for when a reaction is considered present, and similarly non-present, where the thresholds are chosen at values that guarantee significance also after correction for multiple testing, regardless of the p values for other reactions. For a reaction to be considered confidently present, we have chosen that > 98% of bootstrap results should include the reaction, and < 2 % for non-present. The thresholds are purposely selected conservatively to keep the false positives at a low level at the expense of more false negatives. To estimate the statistical significance at these levels, we consider a pair of cell types and apply a Fisher’s exact test with the reaction present in 99 bootstraps for one cell type present in 1 bootstrap for the other, yielding $p < 2.2*10^{-16}$. We can then estimate the minimal number of reactions $n_{min}$ required in the model to render such a p value non-significant after correction for multiple testing using the Benjamini-Hochberg method:

$$n_{min} = \frac{i}{p}Q$$

where *i* is the p-value rank, *p* is the p value, and *Q* is the false positive rate, which we set to 0.05. In the worst-case scenario, where the adjusted p-value will reach its maximum value, the reaction we are investigating will have the lowest p value of all reactions, giving the rank *i*=1. Furthermore, we assume that p= $2.2*10^{-16}$ (and not smaller). Using these values yields $n_{min}$ = $2.3*10^{14}$ reactions. With fewer reactions, which in the metabolic model is on the order of 10,000, it is not possible that a correction for multiple testing will yield an adjusted non-significant p value, and hence, there is no need to perform a pairwise comparison of the p values of cell types when using the thresholds.

### Supplementary note 2 – ftINIT

The purpose of the ftINIT (fast task-driven Integrative Network Inference for Tissues) algorithm is to determine the available metabolic subnetwork in a sample based on a template model and omics data. The ftINIT algorithm is as previous versions based on assigning scores to reactions, in this case from RNA-Seq data and gene rules (GPRs) as described previously (18,19). The previous version of the algorithm was based on two steps: The network minimization step and gap filling based on essential metabolic tasks.

The network minimization step strives to determine which reactions to include in the model by maximizing the sum of the reaction scores of the included reactions, while ensuring that all included reactions can carry flux. The optimization problem in this step is based on mixed integer linear programming (MILP) and is a computationally expensive problem for a large model such as Human-GEM. To speed up the process, the optimization can be run with two flags that allow for secretion of all metabolites and for reversible reactions to carry flux in both directions. In practice, the latter means that all reversible reactions can form loops within themselves and can thereby always carry flux, and any such reactions with positive scores will automatically be included in the resulting model. The major downside of using these flags is that unwanted gaps can be left in the model, but for use with Human1, they are required to reach an acceptable execution time.

In the gap-filling step, the algorithm traverses a list of metabolic tasks, where a certain product(s) should be possible to produce from a given substrate(s). For each task the algorithm ensures that the task can be performed by adding reactions at a minimal cost to the total reaction score of the included reactions.

The purpose of ftINIT is to provide a substantially faster software for generating context-specific models from RNA-Seq data with comparable model output quality. To accomplish this goal, we have applied 3 strategies to the network minimization step: 1) Optimization of the existing code (not further described), 2) Reformulation of the MILP, and 3) A split of the network minimization step into several sub-steps. In addition, the gap-filling code was optimized, reducing the execution time of that step by an order of magnitude (not further described). ftINIT is primarily optimized for use with the Human1 model.

#### Reformulation of the MILP

A key factor for reducing the MILP solve time is to reduce the number of variables in the problem, especially the number of integer (Boolean) variables. In the previous versions of tINIT, the model was converted to an irreversible model, in which each reaction was given a boolean variable describing if the reaction should be included or not in the final model. In ftINIT, we have reduced the number of integer variables using approaches such as: 1) We do not convert the model to an irreversible model, which leads to fewer reactions. It also solves the problem with loops caused by reversible reactions. 2) Irreversible reactions with a positive score do not need an integer variable; the problem can be defined using only continuous variables for these reactions. 3) All linearly dependent reactions are merged (only when running the MILP). 4) Some reversible reactions are made irreversible (such as reactions identified as essential, or reactions that can only carry flux in one direction for topological reasons). 5) Common metabolites, e.g., H+, H2O, etc., are removed from the model, since they are expected to be available and not practically influence the MILP, which reduces the complexity of the problem. Collectively, these optimizations substantially reduce the complexity and the number of integer variables in the MILP.

#### Division of the network minimization step into sub-steps

An obvious method to shorten the execution time of the MILP in the network minimization step of the problem is to reduce the size of the problem. The Human-GEM model contains many reactions without GPRs, which means that RNA-Seq data cannot be used to score these reactions. Such reactions are by default assigned a score of -2, and will in tINIT/ftINIT not be added unless they are needed to provide flux for other reactions with positive scores. In total, there are several thousand reactions without GPRs, and an alternative method to scoring them with a -2 is to not score some of them at all, and thereby do not include them in the optimization, while still allowing them to carry flux. In ftINIT, we have implemented this possibility. To enable the user to select which reactions to omit from the optimization problem, they are divided into 8 partly overlapping categories, and selected individually with a vector of 8 Booleans, for example [1 1 1 1 1 1 1 0], where the reactions selected is the union of the specified categories (Table A). The vector of Booleans is referred to as the 'rxns to ignore mask'.

***Table A. The 8 different categories of reactions without GPRs that can be selected for exclusion from the optimization problem.*** *The number of custom reactions represent the number of such reactions used in this study.*

| Id | Name | Description | No. rxns |
|---|---|---|---|
| 1 | Exchange | All exchange reactions. | 1,110 |
| 2 | Import | Reactions importing metabolites into the cell. | 706 |
| 3 | Simple transport | Reactions that transport one metabolite from one compartment to another. | 821 |
| 4 | Advanced transport | Transport reactions that involve multiple metabolites, e.g., antiporters. | 129 |
| 5 | Spontaneous | Reactions marked as spontaneous in the model. | 10 |
| 6 | EC | Extracellular reactions, i.e. reactions in the s compartment. | 32 |
| 7 | Custom | Reactions defined by the user. | 69 |
| 8 | All | All reactions without GPRs. | 3,466 |

We divided the network minimization step of ftINIT into 3 sub-steps, of which the third is optional (Fig. A). As mentioned above, the previous tINIT algorithm was typically run with flags that allowed for secretion of all metabolites and for reversible reactions to carry flux in both directions. To optimize the code, we therefore used the same simplification for the first sub-step. Since the model is no longer irreversible in ftINIT, the second simplification cannot be implemented, but we instead omit all reversible reactions with positive reaction score from the problem, since they would automatically be turned on in the original version of tINIT. Likewise, the reactions without GPRs as selected by the user are excluded from the problem.

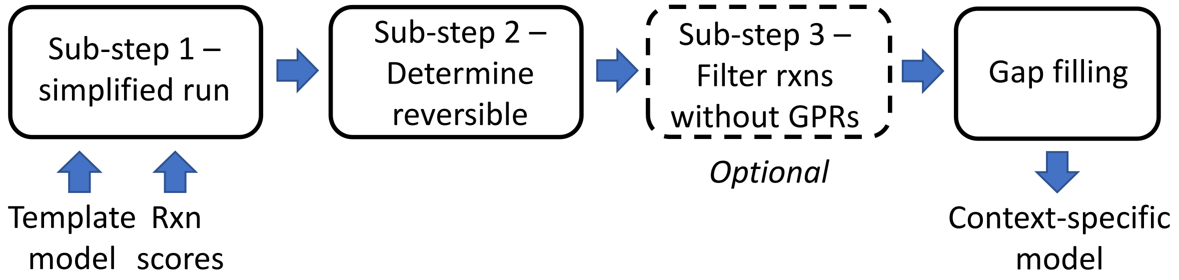


***Fig. A: Sub-steps in the ftINIT algorithm.***

In the second sub-step, we include the reversible reactions with positive reaction scores in the problem, while omitting the reactions decided to be present by the optimization in the first sub-step. This sub-step will thus determine which of the reversible reactions with positive reaction scores that should be added and in addition add reactions not previously added that are needed for those reversible reactions to carry flux.

In the third step, which is optional, we flag all previously included reactions as essential, turn off metabolite secretion, and allow for a different selection of reactions without GPRs that should be excluded from the problem. If for example only the exchange reactions are omitted from the problem in this step, the algorithm will for the rest of the reactions without GPRs determine if they should be included or not.

At the end of the network minimization step, all reactions without GPRs that were not part of the problem in the calculations will simply be included in the model.
